# Hedgehog signaling is required for the maintenance of mesenchymal nephron progenitors

**DOI:** 10.1101/2023.08.12.553098

**Authors:** Eunah Chung, Patrick Deacon, Yueh-Chiang Hu, Hee-Woong Lim, Joo-Seop Park

## Abstract

Mesenchymal nephron progenitors (mNPs) give rise to all nephron tubules in the mammalian kidney. Since premature depletion of these cells leads to low nephron numbers, high blood pressure, and various renal diseases, it is critical that we understand how mNPs are maintained. While Fgf, Bmp, and Wnt signaling pathways are known to be required for the maintenance of these cells, it is unclear if any other signaling pathways also play roles. In this report, we explored the role of Hedgehog signaling in mNPs. We found that loss of either Shh in the collecting duct or Smo from the nephron lineage resulted in premature depletion of mNPs. Transcriptional profiling of mNPs with different Smo dosages suggested that Hedgehog signaling inhibited Notch signaling and upregulated the expression of Fox transcription factors such as *Foxc1* and *Foxp4*. Consistent with these observations, we found that ectopic expression of *Jag1* caused the premature depletion of mNPs as seen in the *Smo* mutant kidney. We also found that Foxc1 was capable of binding to mitotic condensed chromatin, a feature of a mitotic bookmarking factor. Our study demonstrates a previously unappreciated role of Hedgehog signaling in preventing premature depletion of mNPs by repressing Notch signaling and likely by activating the expression of Fox factors.

**TRANSLATIONAL STATEMENT:** Premature depletion of nephron progenitors results in low nephron endowment, leading to high blood pressure and various renal diseases. Sound understanding of the molecular mechanisms underlying the maintenance of nephron progenitors is required for intervention. Although defective Hedgehog signaling is known to cause Pallister-Hall syndrome, its activity in nephron progenitors has been elusive. Here we report that Hedgehog signaling plays an important role in maintaining nephron progenitors. Our findings suggest that Hedgehog signaling pathway is a potential target for enhancing nephron endowment.

## INTRODUCTION

In the developing mammalian kidney, *Six2*-expressing mesenchymal nephron progenitors (mNPs) are multipotent because they have the ability to develop into distinct nephron segments,^1–3^ each responsible for unique physiological functions to serve as blood filtration units. Premature depletion of these cells leads to low nephron numbers, increasing the risk of high blood pressure and various renal diseases.^4^ Hence, it is critical to understand how mNPs are maintained. It is known that Fgf, Bmp, and Wnt signals are required for the maintenance of these cells^5–16^ and this knowledge made it possible to establish *in vitro* culture systems of mNPs.^17–20^ However, it is unclear if any other signaling pathways also play roles in preventing the premature depletion of mNPs.

The Hedgehog signaling pathway plays crucial roles in embryonic development and tissue homeostasis. This pathway can be activated by any of three ligands: Sonic Hedgehog (Shh), Indian Hedgehog (Ihh), and Desert Hedgehog (Dhh). In the absence of the Hedgehog protein, a transmembrane receptor called Patched (Ptch1 and Ptch2) inhibits another transmembrane protein called Smoothened (Smo), preventing the activation of downstream signaling events. The binding of the Hedgehog protein to Ptch relieves its inhibition on Smo, whose activation eventually results in the stabilization of full-length Gli transcription factors (Gli1, Gli2, and Gli3) and Gli-mediated transcriptional activation. *GLI3* mutations producing truncated GLI3 causes renal malformations in human patients with Pallister-Hall syndrome and in a mouse model.^21–23^ While it has been shown that Hedgehog signaling regulates various aspects of kidney development,^24–30^ its role in mNPs has been elusive. Here we show that Hedgehog signaling is required for the maintenance of mNPs. We also show that, in mNPs, Hedgehog signaling represses Notch signaling and upregulates the expression of *Fox* transcription factors including *Foxc1* and *Foxp4*.

## METHODS

### Mice

*Foxc1* conditional knock out (flox) allele (*Foxc1^c^*) was made via a same-day sequential loxP insertion approach. Briefly, the 5’ and 3’ single-guide RNAs (target sequence: GTGGAGAGCAAATGTGATGT and CTCTAAGAGTGCCGGGAATA, respectively) were selected according to the on- and off-target scores from the web tool CRISPOR^31^ and synthesized from Integrated DNA Technologies (IDT). To form the ribonucleoprotein complex (RNP), individual sgRNA (60ng/μl) was mixed with Cas9 protein (IDT; 80ng/μl) in Opti-MEM (ThermoFisher) and incubated at 37°C for 15 minutes. The donor oligo (Ultramer from IDT) with asymmetrical homologous arm design^32^ and the loxP sequence was added to the respective RNP at the final concentration of 500ng/μl. The zygotes from superovulated female mice on the C57BL/6 background were first electroporated with 7μl RNP/donor mix targeting the 5’ site on ice using a Genome Editor electroporator (BEX; 30V, 1ms width, and 5 pulses with 1s interval). Two minutes after electroporation, zygotes were moved into 500μl cold M2 medium (Sigma), warmed up to room temperature, and then cultured in KSOM (CytoSpring) containing 2μM M3814 (AOBIOUS) in a CO_2_ incubator. Four hours later, the zygotes were electroporated again to target the 3’ site and subsequently transferred into the oviductal ampulla of pseudopregnant CD-1 females for birth. The offspring carrying 5’ and 3’ loxP in cis were identified by PCR and Sanger sequencing. *Foxc1^c^* was genotyped with two oligonucleotides (TGGGAATAGTAGCTGTCAGATGGC and TCTGTTCGCTGGTGTGAGGAATC, wild type = 229bp, flox = 269bp). Other mouse alleles used in this study have been previously published: *Rosa26^Sun^*^1,33^ *Shh^Cre^*,^34^ *Calb1^Cre^*,^35^ *Shh^c^*,^36^ *Smo^c^*,^37^ *Six2GFPcre*,^1, 3^ *Smo^-^*,^38^ *Rosa26^SmoM^*^2,39^ *Rosa26^NuTrap^*,^40^ *Rosa26^Jag^*^1^.^41^ All experiments were performed in accordance with animal care guidelines and the protocols were approved by the Institutional Animal Care and Use Committee of Cincinnati Children’s Hospital Medical Center or Northwestern University. We adhere to the NIH Guide for the Care and Use of Laboratory Animals.

### Immunofluorescence staining

Kidneys were fixed in 4% paraformaldehyde in phosphate-buffered saline (PBS) for 15 minutes, incubated overnight in 10% sucrose/PBS at 4°C, and imbedded in OCT (Fisher Scientific). Cryosections (8-9μm) were incubated overnight with primary antibodies in 5% heat-inactivated sheep serum/PBST (PBS with 0.1% Triton X-100). We used primary antibodies for GFP (1:500, Aves GFP-1020;1:900, Synaptic Systems 132 005), Wt1 (1:100, Santa Cruz sc-7385), Cytokeratin (1:200, Sigma C2562), Six2 (1:200, Abcam ab277942;1:500, Novus H00010736-M01), Calb1 (1:500, Abcam ab82812), Slc12a3 (1:300, Sigma HPA028748), Pbx1 (1:500, Proteintech 18204-1-AP), Cdh1 (1:500, Santa Cruz sc-59778), Pax2 (1:200, Proteintech 21385-1-AP), Jag1 (1:20, Developmental Studies Hybridoma Bank TS1.15H), Foxc1 (1:100, Sigma HPA040670), Foxp4 (1:100, Sigma HPA007176), Lhx1 (1:20, Developmental Studies Hybridoma Bank 4F2), and Krt18 (1:50, Santa Cruz sc-32329). Fluorophore-labeled secondary antibodies were used for indirect visualization of the target. Images were taken with a Nikon Ti-2 inverted widefield microscope equipped with an Andor Zyla camera and Lumencor SpectraX light source housed at the Confocal Imaging Core (CIC) at Cincinnati Children’s Hospital Medical Center (CCHMC).

### *In situ* hybridization by RNAscope

To visualize *Six2* or *Jag1* expression, fixed frozen sections of kidneys were processed for RNAscope assay. We used the Six2 probe 500011-C3 and Jag1 probe 412831-C2 for hybridization and developed each channel with fluorescent dyes Opal Reagent 620 or 690 Reagent Pack (Akoya Biosciences) using RNAscope Multiplex Fluorescent Reagent Kit v2 (Advanced Cell Diagnostics, Bio-Techne #323100) following the manufacturer’s instructions. Slides were imaged on a Nikon Ti2 widefield fluorescent microscope with a 10x objective lens.

### Nephron counts

Glomeruli counts were done as previously described.^42^ Briefly, postnatal day 14 kidney was minced with a razor blade and incubated in 2.5ml 6N HCl for 60 minutes followed by repeated pipetting to disrupt further. To the suspension, 10ml of deionized water was added, mixed, and incubated overnight at 4°C. A set volume of macerate (0.1ml to 1ml) was moved to a gridded 6-well dish (Pioneer Scientific PSMP6G) and the experimenter scored the glomeruli blind to the genotype.

### Nephron progenitor cell counts

Quantification of nephron progenitor cells were done as previously described.^29^ From immunostained kidney sections, the number of Six2+ cells per collecting duct tip was scored. Six to seven fields were counted per kidney from middle sections. Two separate Smo loss-of-function (LOF) litters were examined. For each litter, two mutant kidneys of *Smo^c/-^;Six2GFPcre* and two control kidneys of *Smo^c/+^;Six2GFPcre* were used. A total of 149 bud tips each were scored from control and mutant. Likewise, two separate Smo gain-of-function (GOF) litters were examined. A total of 64 bud tips from the mutant kidneys (*Rosa26 ^SmoM^*^2^*^/+^;Six2GFPcre*) and 56 bud tips from the control kidney (*Rosa26^SmoM^*^2^*^/+^*) were scored. Data were shown as the number of Six2+ cells per bud tip. Student’s paired t-Test with a two-tailed distribution was used to calculate *p* values.

### Fluorescence activated cell sorting (FACS) and RNA-seq

Six2GFPcre+ embryonic kidneys were dissociated using TrypLE Select Enzyme 10X (Gibco) and mechanical disruption by repetitive pipetting. Cells were washed in PBS containing 10% fetal bovine serum (FBS), then resuspended in PBS containing 1% FBS and 10mM EDTA and resuspended cells were filtered through 40μm Nylon Cell Strainer (BD Falcon) and kept on ice until FACS. The GFP+ cells were isolated using BD FACSAria II or BD FACSAria Fusion (BD Biosciences) device at the Research Flow Cytometry Core at Cincinnati Children’s Hospital. Collected GFP+ cells were spun down and cell pellet was stored in -80°C until RNA extraction. Bulk RNA-seq with two samples for each genotype was performed as previously described.^43^ Total RNA from FACS isolated cells were prepared using Single Cell RNA Purification Kit (Norgen). From total RNA, mRNA was isolated using NEBNext Poly(A) mRNA Magnetic Isolation Module (E7490, New England Biolabs) following manufacturer’s instructions. Fragmentation of RNA followed by reverse transcription and second strand cDNA synthesis was done using NEBNext RNA First Strand Synthesis Module (E7525L) and NEBNext RNA Second Strand Synthesis Module (E6111L) following manufacturer’s instructions. The resulting double-stranded cDNA’s were further processed to DNA sequencing libraries using ThruPLEX DNA-seq 12S Kit (R400428, Clontech Laboratories). Libraries were size-selected by gel purification for an average size of 350bp. Each purified library was quantified using a Qubit fluorometer (Life Technologies) and equal amounts of each library were pooled and submitted for sequencing on the Illumina NextSeq 500 by the DNA Sequencing and Genotyping Core at CCHMC. All RNA-seq reads were aligned to UCSC mouse genome 10 mm using STAR aligner.^44^ Only uniquely aligned reads were used for downstream analysis. Raw read counts for each gene were measured using FeatureCounts in the subread package with an option, “-s 2 -O --fracOverlap 0.8”.^45^ Differential gene expression analysis was performed using edgeR.^46^ Genes with fold-change > 1.5 and FDR < 0.05 were selected as differentially expressed genes.

Hierarchical clustering was performed using Pearson correlation coefficient as a similarity measure under Ward’s criterion. Linear regression analysis of the gene copy was performed using R, and p-values were adjusted to FDR to identify significantly correlated genes. Data were deposited to Gene Expression Omnibus (GSE237179). Gene ontology (GO) analysis was performed using Enrichr^47–49^ on Groups III and VIII genes. Cell cycle analysis was performed using the gene sets defined in a previous report.^50^ RPKM-normalized gene expression levels were log-normalized and transformed to Z-score for heatmap visualization.

### Transcription factor binding on mitotic chromosomes

HEK293A cells were plated on a chambered coverslip (μ-Slide 8 Well Glass Bottom, ibidi #80827) and transfected the next day with 0.8μg plasmid DNA expressing GFP-tagged Six2 (p1054), FoxA1 (p1031),^51^ or Foxc1 (p1104) using 2μl Lipofectamine 2000. The next day, transfection media was replaced with fresh media containing nocodazole at 330nM final concentration. We set up confocal imaging to start at 15 hours after nocodazole treatment and finish by 18 hours. Hoechst 33342 was added just before microscopy. We imaged live cells on a Nikon A1R inverted LUNV confocal laser scanning microscope equipped with a heated stage insert providing CO_2_ and humidity (Tokai Hit incubator) housed at the Confocal Imaging Core (CIC) at CCHMC. We looked for condensed mitotic chromosomes and captured those images with a 60x oil-immersion objective lens and determined if GFP fusion proteins were colocalized.

## RESULTS

### Loss of Shh causes renal hypoplasia

It was reported that, in the developing mouse kidney, *Shh* is expressed in the medullary collecting ducts and the ureter.^24, 52^ Consistent with this, we found that *Shh^Cre^*, whose Cre expression is controlled by the *Shh* locus, activated the *Rosa26^Sun^*^1^ reporter mainly in the medullary collecting ducts and the ureter (Figure 1A). To investigate the roles of Shh in kidney development, we conditionally deleted *Shh* from the kidney using *Calb1^Cre^*.^35^ Similar to *Hoxb7^cre^*,^24^ *Calb1^Cre^* mainly targets the collecting duct and the ureter in the kidney (Supplemental Figure 1). The *Shh* mutant kidney was considerably smaller than the control kidney (Figure 1B). Six2 immunostaining showed that Six2+ mNPs were noticeably missing at multiple positions at the cortex of the mutant kidney (Figure 1C, while arrowheads), raising an interesting possibility that Hedgehog signaling is required for the maintenance of mNPs. It has been reported that deletion of *Smo* in the interstitial lineage (*Smo^c/-^;Foxd1^cre/+^*) results in severe patterning defects in the nephrogenic zone, suggesting that Hedgehog signaling is active in the cortical stroma.^29^ We found that the *Shh* mutant kidney showed gaps in Pbx1+ cortical interstitial cells (Supplemental Figure 2), resembling the *Smo^c/-^;Foxd1^cre/+^* mutant kidney, which suggested that Shh is the ligand signaling in the cortical interstitial cells. While it was reported that the nephron progenitor domain was expanded in *Smo^c/-^;Foxd1^cre/+^* mutant kidney,^29^ we found that, in the *Shh* mutant kidney, many of the ureteric tips had few or no Six2+ cells (Figure 1C). These results led us to investigate the role of Hedgehog signaling in mNPs.

**Figure 1.**
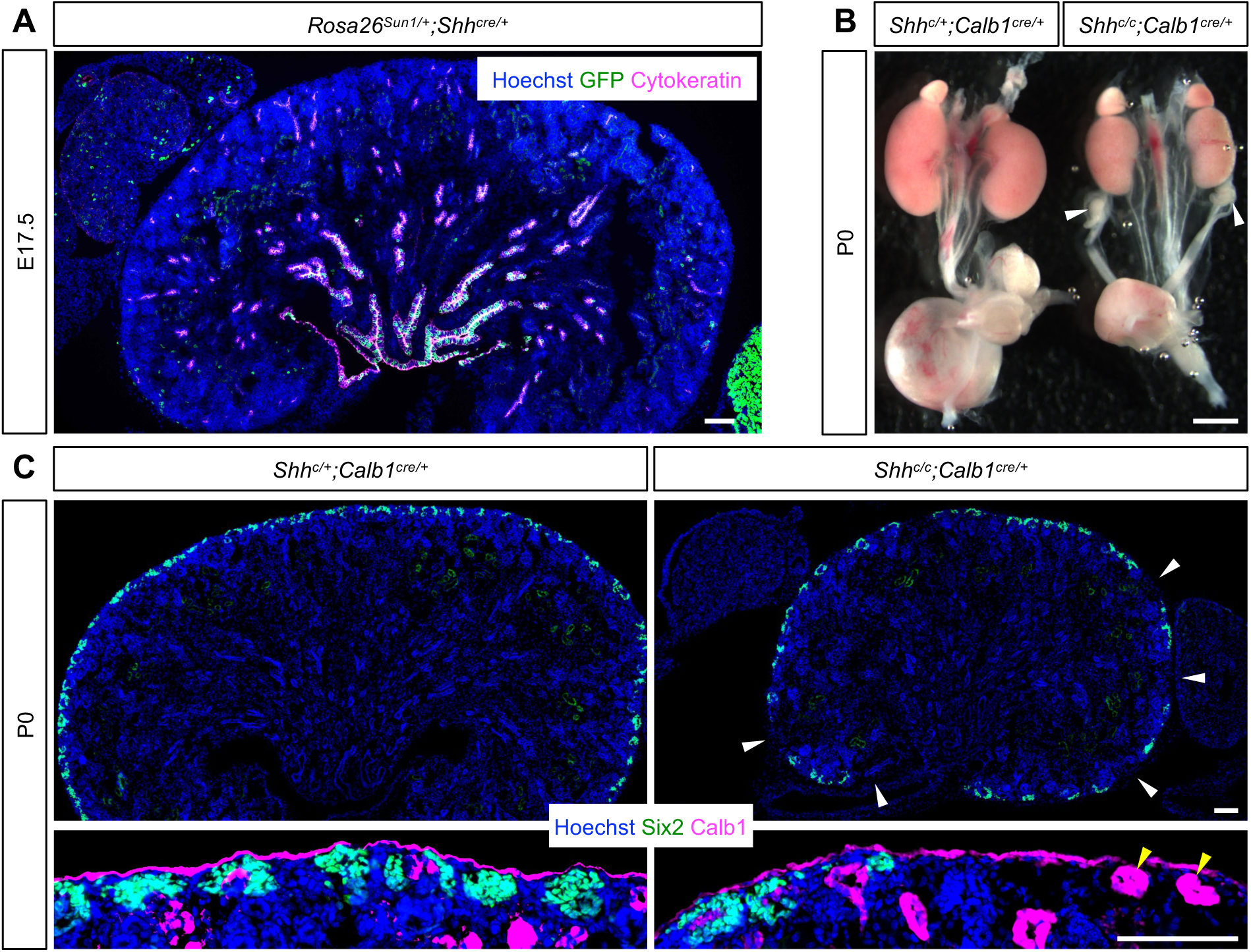
*Shh* is expressed in the medullary collecting duct in the developing mouse kidney and its deletion causes renal hypoplasia accompanied by loss of Six2+ nephron progenitors. (A) Lineage analysis of Shh+ cells in the mouse kidney at E17.5. *ShhCre* activates the Rosa26 Sun1 reporter mainly at the medullary collecting duct. Scale bar, 100μm. (B) Conditional deletion of *Shh* from the collecting duct with *Calb1Cre*. The *Shh* mutant kidney is smaller than the control kidney at P0. Scale bar, 1mm. (C) In the newborn control kidney, Six2+ cells are present throughout the cortex of the kidney. The cortex of the *Shh* mutant kidney has several regions, where Six2+ cells are absent (white arrowheads). Yellow arrowheads mark ureteric tips without adjacent Six2+ cells. Scale bar, 100μm. Image is representative of *n*=3.

### Loss of Smo in the nephron lineage causes premature depletion of mNPs

In order to test if Hedgehog signaling is required for the maintenance of mNPs, we conditionally deleted *Smo*, the gene encoding an essential component of Hedgehog signaling (Smoothened), using transgenic *Six2GFPcre*, which targets mNPs and their descendants.^1, 3^ *Six2GFPcre*-mediated deletion of *Smo* would inhibit Hedgehog signaling in the entire nephron lineage including mNPs. Similar to the *Shh* mutant kidney, we found that the *Smo* mutant kidney was considerably smaller than the control kidney with weaker *Six2GFPcre* expression (Figures 2A & 2B). Six2 immunostaining showed that fewer Six2+ mNPs were present in the *Smo* mutant embryonic kidneys by *Six2GFPcre* (Figure 2C and Supplemental Figure 3). We found that the *Smo* mutant kidney was still smaller than the control kidney at 2 weeks with almost 50% reduction in nephron number (Figures 2D & 2E). Unlike the *Shh* mutant kidney, the *Smo* mutant kidney did not show a gap of Pbx1+ cells (Supplemental Figure 2), consistent with the notion that blockage of Hedgehog signaling is specific to the nephron lineage. Taken together, these results show that loss of Smo causes premature depletion of mNPs and reduced nephron endowment, supporting the idea that Hedgehog signaling is required for the maintenance of mNPs.

**Figure 2.**
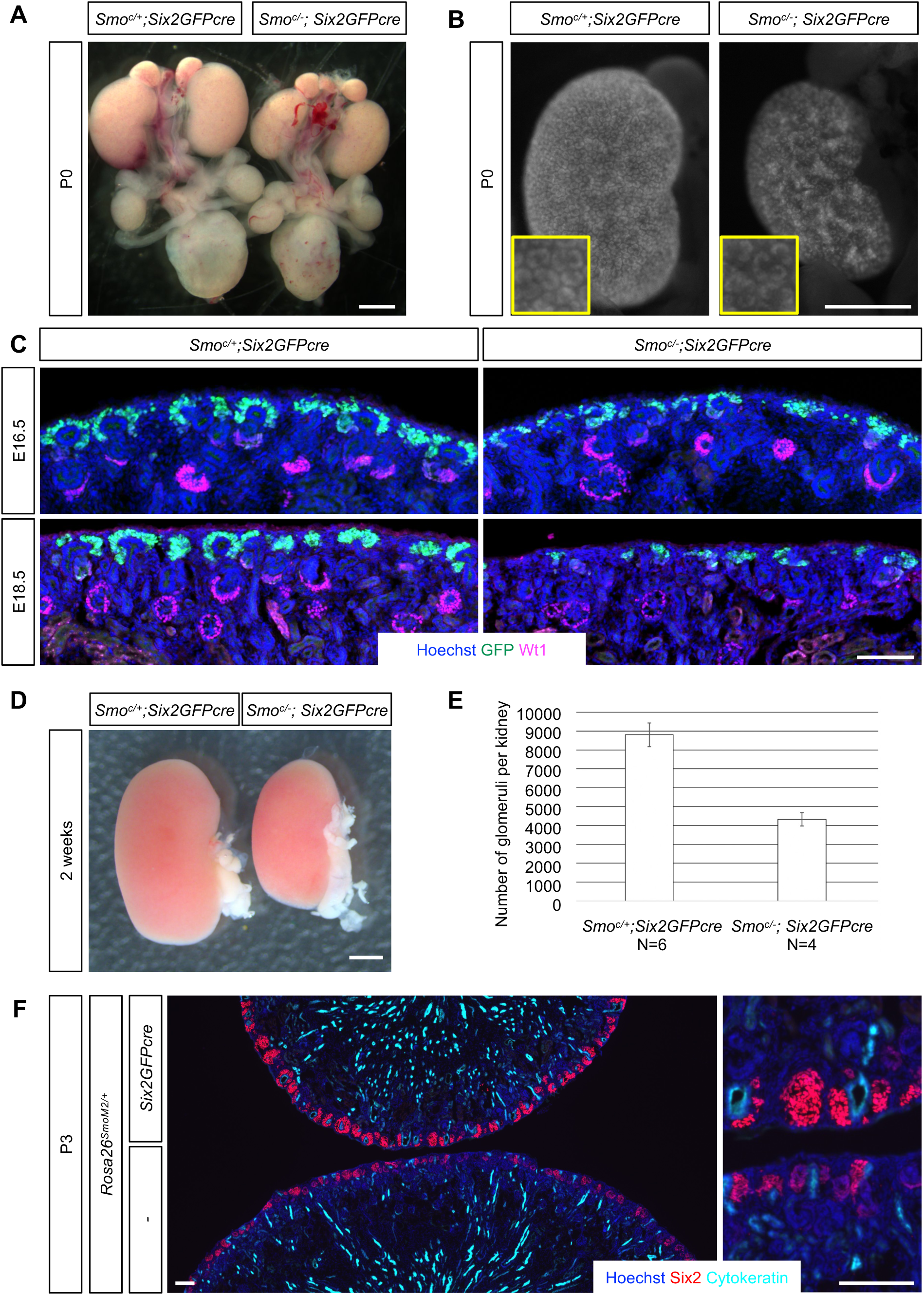
Conditional deletion of *Smo* with *Six2GFPcre* leads to loss of Six2+ nephron progenitors. (A) Conditional deletion of *Smo* from the nephron lineage with *Six2GFPcre*. The *Smo* mutant kidney is smaller than the control kidney at P0. Scale bar, 1mm. (B) *Six2GFPcre* expression is weaker in the *Smo* mutant kidney at P0. Scale bar, 1mm. (C) Fewer Six2+ cells are present in the *Smo* mutant kidney at E16.5 and E18.5. Scale bar, 100μm. (D) The *Smo* mutant kidney is smaller at 2 weeks. Scale bar, 1mm. (E) Nephron endowment is reduced by about 50% in the *Smo* mutant kidney. *p* = 0.00320 (Student’s paired t-Test with a two-tailed distribution.). (F) While the *SmoM2* gain-of-function mutant is smaller than the control kidney, the mutant kidney has more Six2+ cells with stronger Six2 signal. Stage, P3. Scale bar, 100μm. Images are representative of *n*=3.

To examine how increased Hedgehog signaling affects mNPs, we carried out a gain-of-function study of *Smo* employing *Six2GFPcre* and *Rosa26-SmoM2* allele, which expresses a constitutively active form of Smoothened upon Cre-mediated recombination.^39^ To test if *SmoM2* expression leads to the persistent presence of mNPs by inhibiting the depletion of these cells, we attempted to examine Six2+ mNPs at P4 when mNPs are normally depleted. However, we were unable to obtain live *SmoM2* mutant pups beyond P3 due to postnatal lethality. We found that *Six2GFPcre*-mediated activation of *SmoM2* caused the mutant kidney to be smaller than the control kidney (Figure 2F). This is likely due to the fact that the presence of *Six2GFPcre* itself makes the kidneys smaller.^1, 53, 54^ Interestingly, while the control kidneys apparently started to show signs of loss of mNPs at P3, the kidneys from the littermate *SmoM2* mutant pups had more Six2+ mNPs with stronger Six2 signal (Figure 2F and Supplemental Figure 3). We found that expression of SmoM2 itself did not block nephrogenesis (Supplemental Figure 4). These results suggest that constitutive activation of Hedgehog signaling inhibits the depletion of mNPs.

### Identification of Hedgehog-responsive genes in mNPs

In order to determine which genes are regulated by Hedgehog signaling in mNPs, we FACS-isolated Six2GFPcre+ mNPs from both loss- and gain-of-function mutants of *Smo* and their control kidneys at E18.5 (Figure 3A), and performed RNA-seq analyses. Since both loss- and gain-of-function mutants have their own corresponding controls, our datasets contain transcriptional profiling data from mNPs with four different doses of *Smo* (Figure 3B). While unsupervised clustering of transcriptome identified eight groups of differentially expressed genes, we focused on Groups III and VIII, which are upregulated and downregulated by Hedgehog signaling, respectively (Figure 3B and Supplemental Table 1). Gene ontology (GO) analyses showed that regulation of transcription (Group III) and mitochondrial ATP synthesis (Group VIII) were the top enriched biological processes (Supplemental Figure 5 and Supplemental Tables 2 & 3). It has been shown that, when mNPs undergo differentiation, a metabolic switch from glycolysis to mitochondrial respiration occurs.^55, 56^ *Ptch1*, the gene encoding a critical component of Hedgehog signaling, is one of the best-known Hedgehog-responsive genes.^57, 58^ We found that there was a striking linear correlation between *Ptch1* expression and *Smo* dosage (Figure 3C). Similarly to *Ptch1*, we found that Hedgehog signaling upregulated the genes encoding Fox transcription factors such as *Foxc1*, *Foxd2*, and *Foxp4* in mNPs (Figure 3C) and that those genes belonged to the top category (regulation of transcription) in GO analysis (Supplemental Figure 5 and Supplemental Table 2).

**Figure 3.**
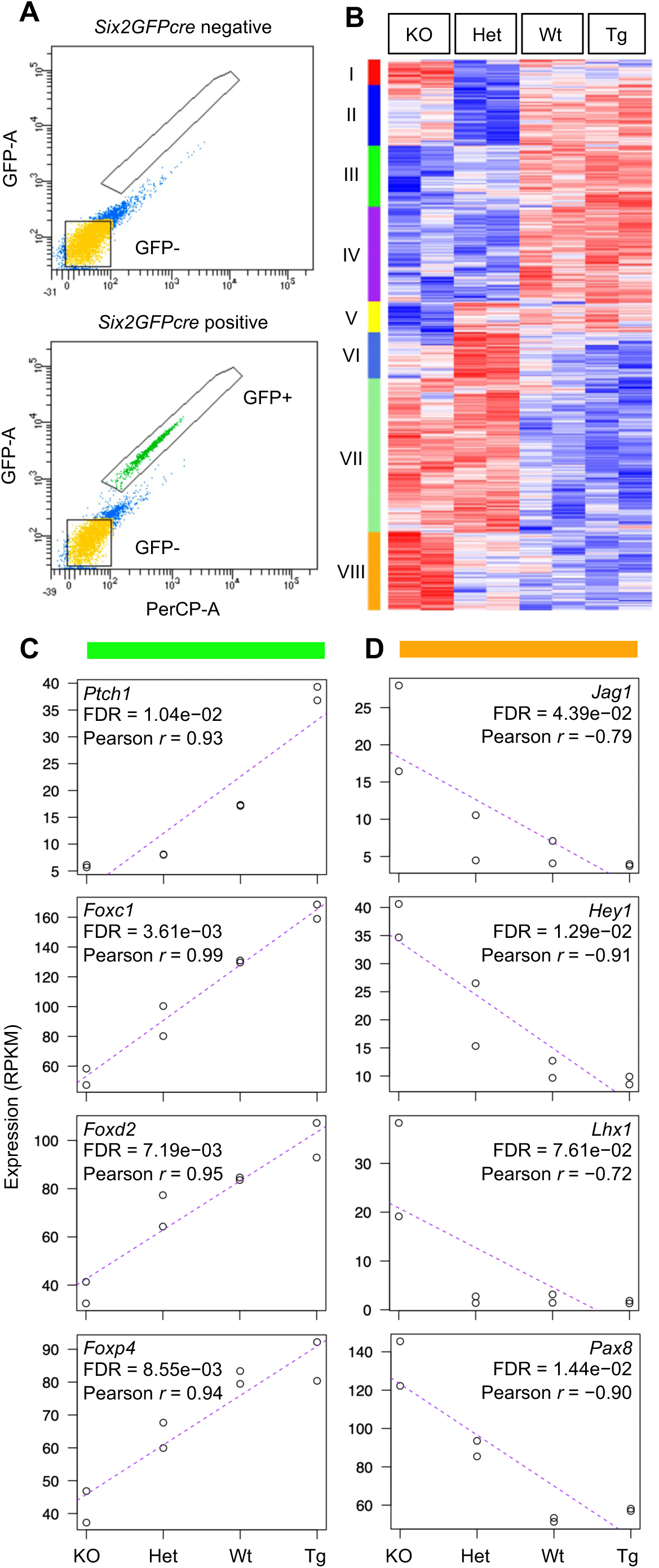
Identification of genes that respond to Hedgehog signaling in the nephron progenitors. (A) FACS isolation of Six2GFPcre+ nephron progenitors (upper, negative control with no GFP; lower, Six2GFPcre+ cells shown in green). (B) Unsupervised clustering of the transcriptome of Six2GFPcre+ nephron progenitors with four different *Smo* gene doses. (KO=*Smo^c/-^;Six2GFPcre*, Het=*Smo^c/+^;Six2GFPcre*, Wt=*Smo^+/+^;Rosa26^+/+^;Six2GFPcre*, Tg=*Smo^+/+^;Rosa26^SmoM^*^2^*^/+^;Six2GFPcre*). KO and Het have zero and one copy of *Smo*, respectively. While Wt has two copies of *Smo*, Tg has one copy of transgenic *SmoM2* in addition to two copies of *Smo*. Group III (marked by the dark green bar) represents genes exhibiting upregulated expression in response to Hedgehog signaling, while Group VIII (marked by the orange bar) represents genes displaying downregulated expression in response to Hedgehog signaling. (C) Representative genes from Group III. (D) Representative genes from Group VIII (C-D). X-axis represents *Smo* gene doses and Y-axis represents RPKM values.

Our GO analysis with Group VIII genes also identified Notch signaling pathway (Supplemental Table 3) and our RNA-seq data showed that *Jag1*, *Hey1*, *Lhx1*, and *Pax8* were upregulated by loss of *Smo* (Figure 3D). *Jag1* encodes the major Notch ligand in the nephron lineage^59^ and *Hey1* is a well-known Notch target gene.^60^ *Lhx1* is also likely a Notch target gene because constitutive activation of Notch signaling causes ectopic expression of *Lhx1* in mNPs.^61^ Microarray analysis from mNPs showed that *Pax8* activation was accompanied by activation of *Hey1* and *Lhx1*.^10^ Given that Notch is the major signal that leads to the differentiation of mNPs,^2, 61, 62^, this result raised an interesting possibility that Hedgehog signaling maintains mNPs by repressing Notch signaling.

We validated our RNA-seq data by immunostaining and RNAscope *in situ* hybridization. Consistent with the RNA-seq data, immunostaining showed that Foxc1 and Foxp4 were positively regulated by Hedgehog signaling in mNPs (Supplemental Figures 6 and 7). When we examined potential activation of Notch signaling in the *Smo* loss-of-function mutant kidney, we found that *Jag1* mRNA activation was more robust where *Six2* mRNA is downregulated (Supplemental Figure 8), suggesting that loss of Smo causes mNPs to become prone to differentiation via Notch pathway.

### Roles of Hedgehog responsive genes in nephron progenitors

A previous report suggested that Foxc1 and Foxc2 are required for the maintenance of mNPs.^63, 64^ Since our RNA-seq data suggest that *Foxc1* was upregulated by Hedgehog signaling while *Foxc2* was not (Figure 3C and Supplemental Table 1), we tested how loss of *Foxc1* alone would affect mNPs. We found that the *Foxc1* mutant kidney by *Six2GFPcre* showed weaker *Six2GFPcre* expression at P2 (Figure 4A), suggesting that fewer mNPs are present in the absence of Foxc1.

**Figure 4.**
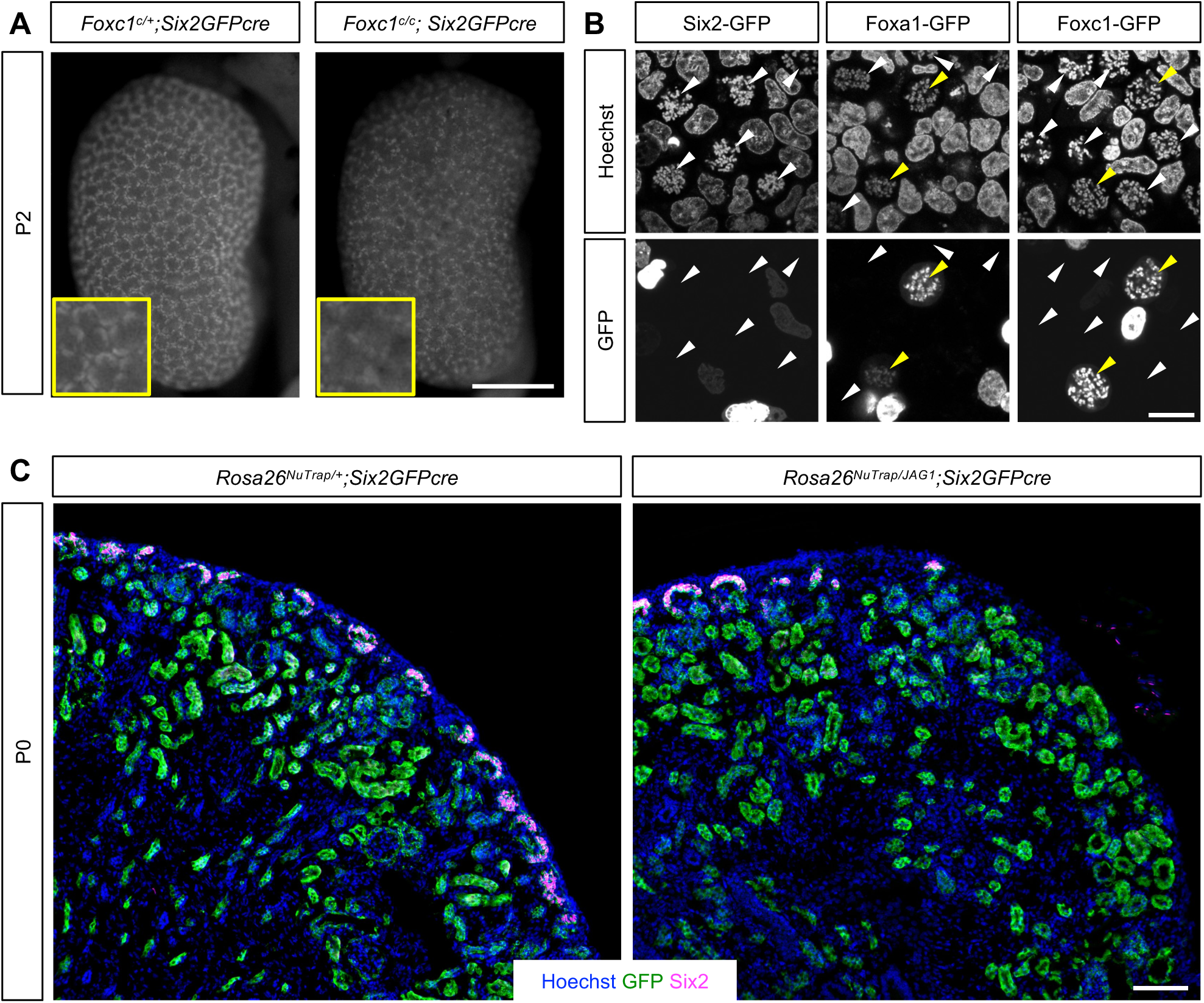
Roles of Hedgehog responsive genes in nephron progenitors. (A) The *Foxc1* mutant kidney shows weaker *Six2GFPcre* expression at P2. Scale bar, 1mm. (B) Foxc1 protein binds to mitotic chromosomes. After transfection and nocodazole treatment on HEK293A cells, we looked for condensed mitotic chromosomes and asked if GFP fusion proteins were colocalized. Foxa1-GFP fusion protein (positive control) was found on mitotic chromatin while Six2-GFP fusion protein (negative control) was never detected on mitotic chromatin and only found in interphase nuclei. Similar to Foxa1-GFP, Foxc1-GFP fusion protein was easily detected on mitotic chromatin. White arrowheads mark mitotic chromatin and yellow arrowheads mark GFP on mitotic chromatin. Scale bar, 10μm. (C) Ectopic expression of *Jag1* with *Six2GFPcre* causes premature depletion of Six2+ nephron progenitors. The Rosa26 NuTrap reporter serves as a lineage tracer for *Six2GFPcre*. Scale bar, 100μm

During mitosis, gene transcription is silenced because most transcription factors dissociate from the condensed chromatin.^65, 66^ However, some transcription factors such as Foxa1^51^ and Hnf1b^67^ can bind to mitotic condensed chromatin. These transcription factors have the potential to “bookmark” their target genes, allowing rapid reactivation after mitosis. Interestingly, we found that Foxc1 was capable of binding to mitotic chromatin (Figure 4B), suggesting that Foxc transcription factors may act as mitotic bookmarking transcription factors in kidney development.

Since our RNA-seq data suggested that loss of Smo caused the upregulation of *Jag1*, we tested if ectopic expression of *Jag1* would phenocopy the *Smo* loss-of-function mutant kidney by employing *Six2GFPcre* and *Rosa26-Jag1* allele, which expresses *Jag1* upon Cre-mediated recombination. We discovered that the *Jag1* gain-of-function mutant kidney had considerably fewer Six2+ mNPs, mimicking loss of Smo (Figure 4C). This result is consistent with previous reports that Notch signaling triggers the differentiation of mNPs.^2, 61, 68^

Taken together, our results suggest that Hedgehog signaling prevents premature depletion of mNPs and that it activates the expression of Fox factors and represses Notch signaling in mNPs.

## DISCUSSION

It was reported that forced expression of a truncated form of Gli3 with *Six2GFPcre* causes premature depletion of mNPs.^28^ While this result was consistent with the fact that the truncation of GLI3 causes renal malformations in human patients with Pallister-Hall syndrome, the phenotype was thought to be caused by a ligand-independent action of the truncated GLI3 and not considered as evidence for active Hedgehog signaling in mNPs.^69^ Our loss-of-function studies of *Shh* and *Smo* mutant kidneys strongly suggest that Hedgehog signaling is active in the cortex of the developing kidney including the Six2+ mNPs and that Shh acts as a ligand, although we cannot rule out the possibility that Shh may act redundantly with Ihh which is expressed in straight proximal tubules.^52^ Consistent with the notion that Hedgehog signaling is required for the maintenance of mNPs, forced expression of *SmoM2* inhibited depletion of mNPs (Figure 2F) without blocking nephrogenesis (Supplemental Figure 4). Further investigation will be required to determine how SmoM2 expression prolongs nephrogenesis.

It has been shown that Hedgehog ligands can act as morphogens by traveling long distances. Possible dispersion mechanisms for morphogens include free diffusion, facilitated diffusion (positive diffusion regulators acting as shuttles), transcytosis (morphogens passing through cells via repeated endocytosis and exocytosis) and cytonemes (long extended filopodia-like cellular structures delivering morphogens by direct contact).^70^ Though it is unclear how Shh from the medullary collecting duct could reach the cortex to regulate mNPs, the distance for Shh to travel to reach mNPs would increase over time during development. Hence, it is likely that mNPs experience dynamic changes in Hedgehog activity at different developmental stages.

Our transcriptome analysis suggests that Hedgehog signaling upregulates *Fox* genes and inhibits Notch signaling in mNPs (Supplemental Figure 9). It has been shown that Hedgehog signaling regulates expression of *Fox* genes in diverse biological contexts.^39, 57, 58, 71–73^ Interestingly, it has recently been shown that deletion of *Foxc1* and *Foxc2* with *Six2CreER* causes the upregulation of *Pax8* and *Hey1* in Six2+ cells,^63^ similar to mNPs lacking *Smo*. Our discovery that Foxc1 can bind to mitotic chromatin suggests that Foxc1, and potentially other Fox factors, may be able to act as a mitotic bookmarking factor in the developing kidney. Our finding that forced expression of *JAG1* phenocopies loss of *Smo* is consistent with previous reports that Notch activation causes premature depletion of mNPs.^2, 68^ Ectopic Notch activation in mNPs may be responsible for cessation of nephrogenesis and inhibition of Notch signaling may be an underlying mechanism for how *SmoM2* expression prolongs nephrogenesis.

In summary, our study demonstrates a previously unappreciated role of Hedgehog signaling in preventing premature depletion of mNPs. In conjunction with other signaling pathways known to be required for mNP maintenance, such as Fgf and Bmp, employing Hedgehog signaling pathway is likely to improve the ability to promote self-renewal of mNPs *in vitro* without losing multipotency.

## DISCLOSURE

All authors declare no competing interests.

## DATA STATEMENT

RNA-seq data were deposited to Gene Expression Omnibus (GSE237179).

## ACKNOWLEDGMENTS

We thank the Confocal Imaging Core (CIC), Research Flow Cytometry Core (RFCC) and the DNA Sequencing and Genotyping Core (DSGC) at CCHMC. Work in J.P.’s laboratory was supported by the National Institute of Diabetes and Digestive and Kidney Diseases, National Institutes of Health (DK120847, DK125577, DK131052, DK127634, and DK120842).

## AUTHOR CONTRIBUTIONS

E.C. performed most of the experiments. P.D. performed glomerular counting. Y.C.H. generated *Foxc1* flox mice. H.W.L. performed bioinformatic analyses. E.C. and J.P. designed the experiments, analyzed the data, and co-wrote the manuscript. J.P. made the figures. All authors approved the final version of the manuscript.

