## Supplemental Figures for "Hedgehog signaling is required for the maintenance of mesenchymal nephron progenitors"

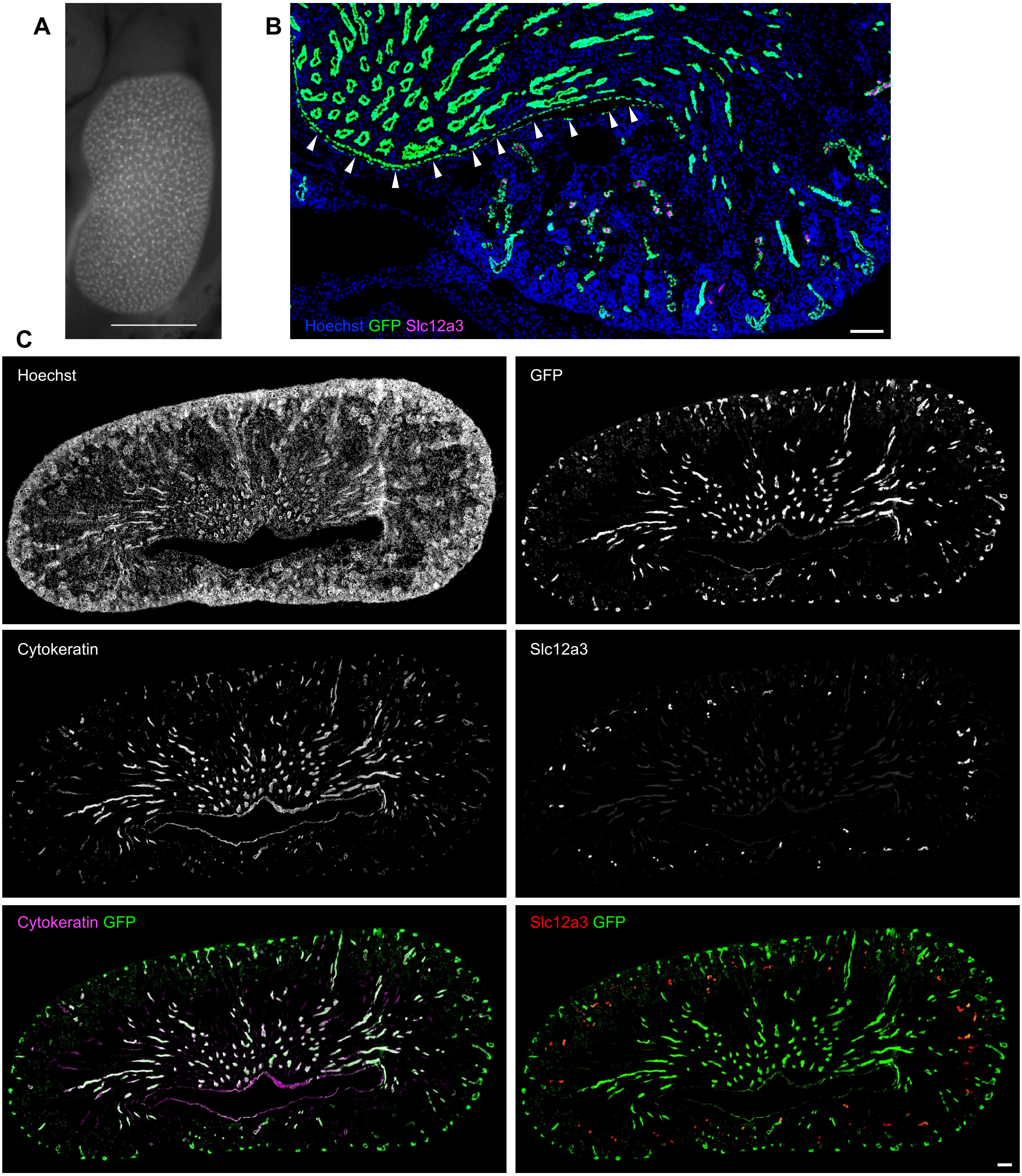


Supplemental Figure 1
(A) EYFP expression pattern of the *Rosa26*$^{Ai3}$ reporter activated by *Calb1*$^{Cre}$ shows that Calb1Cre targets the collecting duct and the ureter. Stage, E18.5. Scale bar, 1mm. (B) *Rosa26*$^{Sun1}$ reporter activated by *Calb1*$^{Cre}$ marks the ureter (white arrowheads) and a subset of Slc12a3+ distal tubules. Stage, P0. Scale bar, 100μm. (C) The *Rosa26*$^{EYFP}$ reporter activated by *Calb1*$^{Cre}$ largely overlaps with cytokeratin+ collecting duct. A subset of Slc12a3+ distal tubules are also positive for GFP. Medullary collecting duct shows stronger cytokeratin staining than cortical collecting duct. Stage, P0. Scale bar, 100μm.

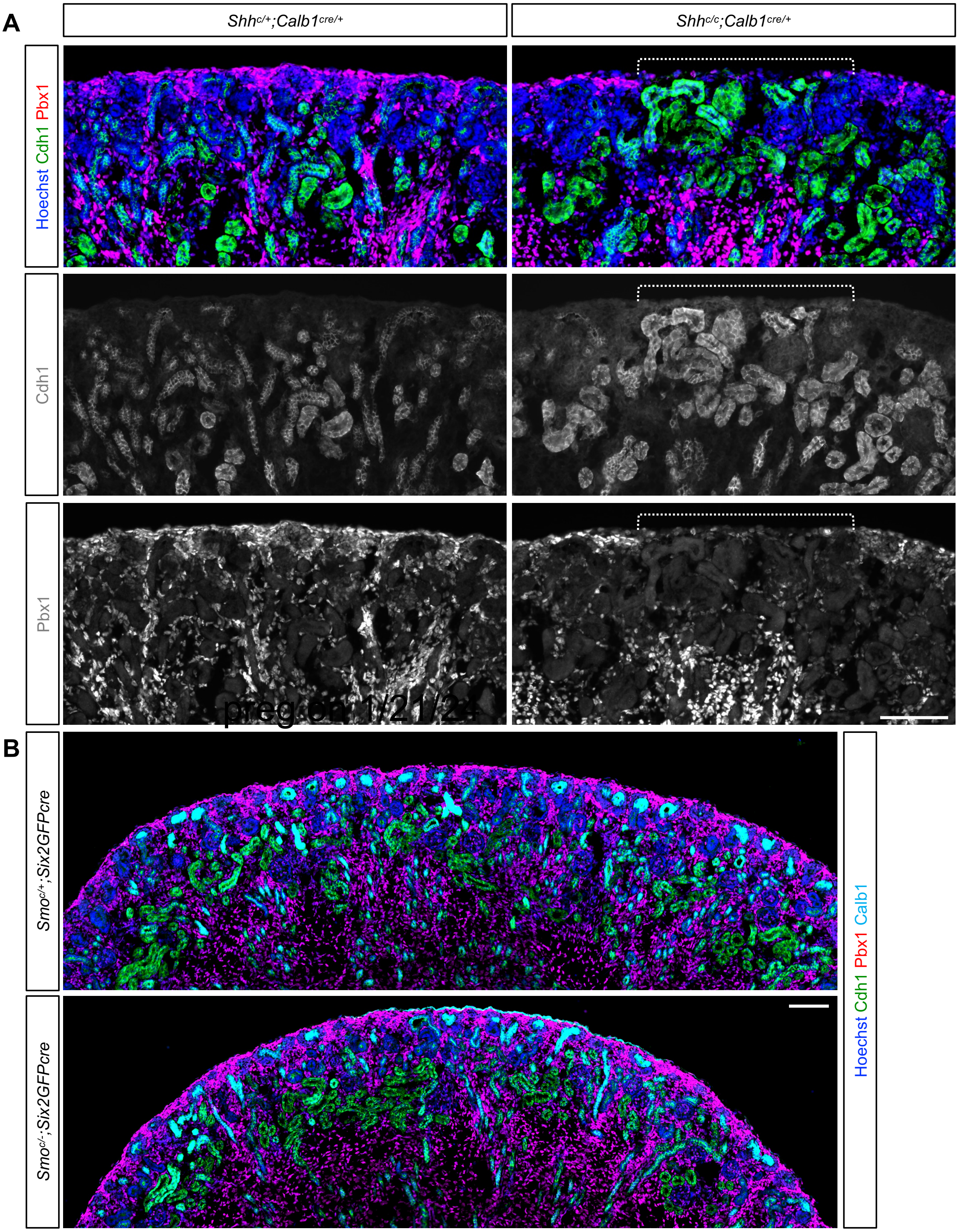


Supplemental Figure 2
Strong Pbx1 signal marks interstitial cells including cortical interstitial progenitors while weak Pbx1 signal marks mesenchymal nephron progenitors. Cdh1 marks nephron tubules and collecting duct. Calb1 marks the collecting duct. (A) Control kidney shows the presence of Pbx1+ cortical interstitial progenitors throughout the entire cortex. In contrast, the *Shh* mutant kidney shows cortical mispatterning. A part of the cortex (dotted bracket) lacks Pbx1+ cortical interstitial progenitors and this area is occupied by ectopic Cdh1+ nephron tubules. Stage, P0. Scale bar, 100μm. (B) *Smo* mutant kidney does not show gaps in Pbx1+ cortical interstitial progenitors. Stage, E18.5. Scale bar, 100μm.

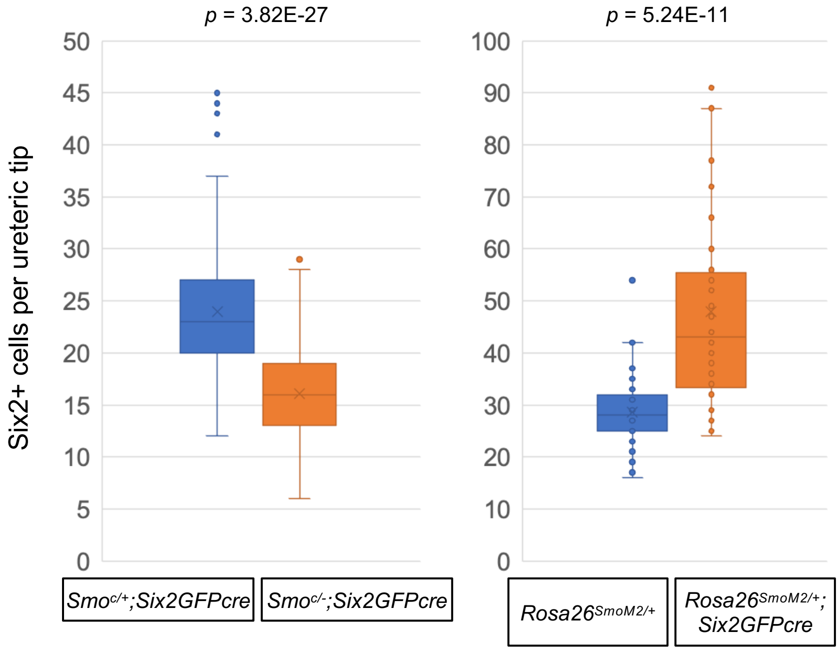


Supplemental Figure 3
Quantitation of Six2+ nephron progenitor cells per bud tip

Left panel: Smo loss-of-function mutant kidneys by Six2GFPcre have fewer Six2+ nephron progenitor cells in the nephrogenic zone. Average number of Six2+ cells/tip in control=23.98 (n=4 kidneys, 149 tips counted). Average number of Six2+ cells/tip in mutant=16.05 (n=4 kidneys, 149 tips counted). Stage E18.5

Right panel: SmoM2 gain-of-function mutant kidneys have more Six2+ nephron progenitor cells. Average number of Six2+ cells/tip in control=28.64 (n=3 kidneys, 64 tips counted). Average number of Six2+ cells/tip in mutant=47.94 (n=4 kidneys, 56 tips counted). Stage P3

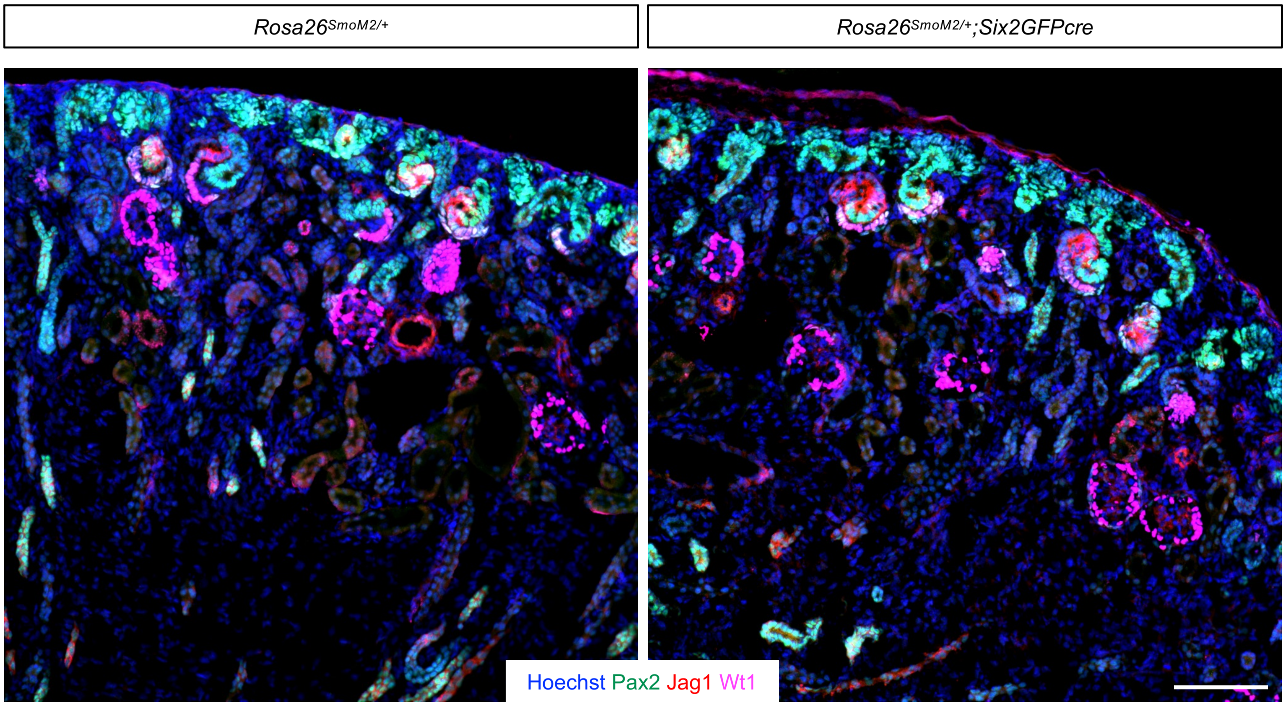


Supplemental Figure 4
Expression of SmoM2 does not block nephrogenesis. Both control and mutant kidneys show active nephrogenesis. Pax2 marks collecting duct, mNPs, and early developing nephrons such as renal vesicles and S-shaped bodies. Jag1 marks median segment of S-shaped bodies while Wt1 marks developing podocytes. Stage, E18.5. Scale bar, 100μm.

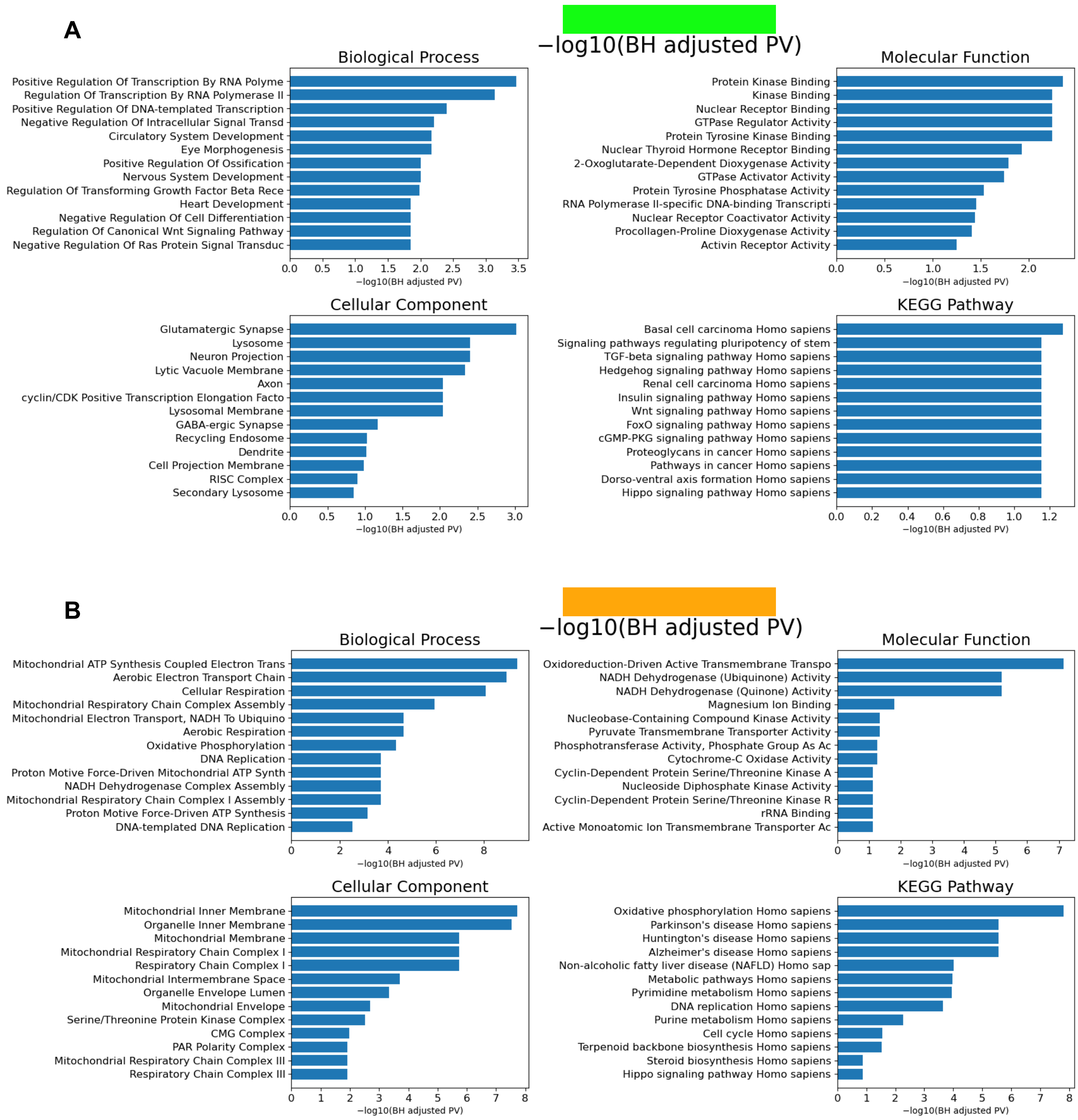


Supplemental Figure 5

Gene ontology analysis with Group III genes (A) and Group VIII genes (B). Full datasets are shown in Supplemental Tables 2 (Group III) and 3 (Group VIII).

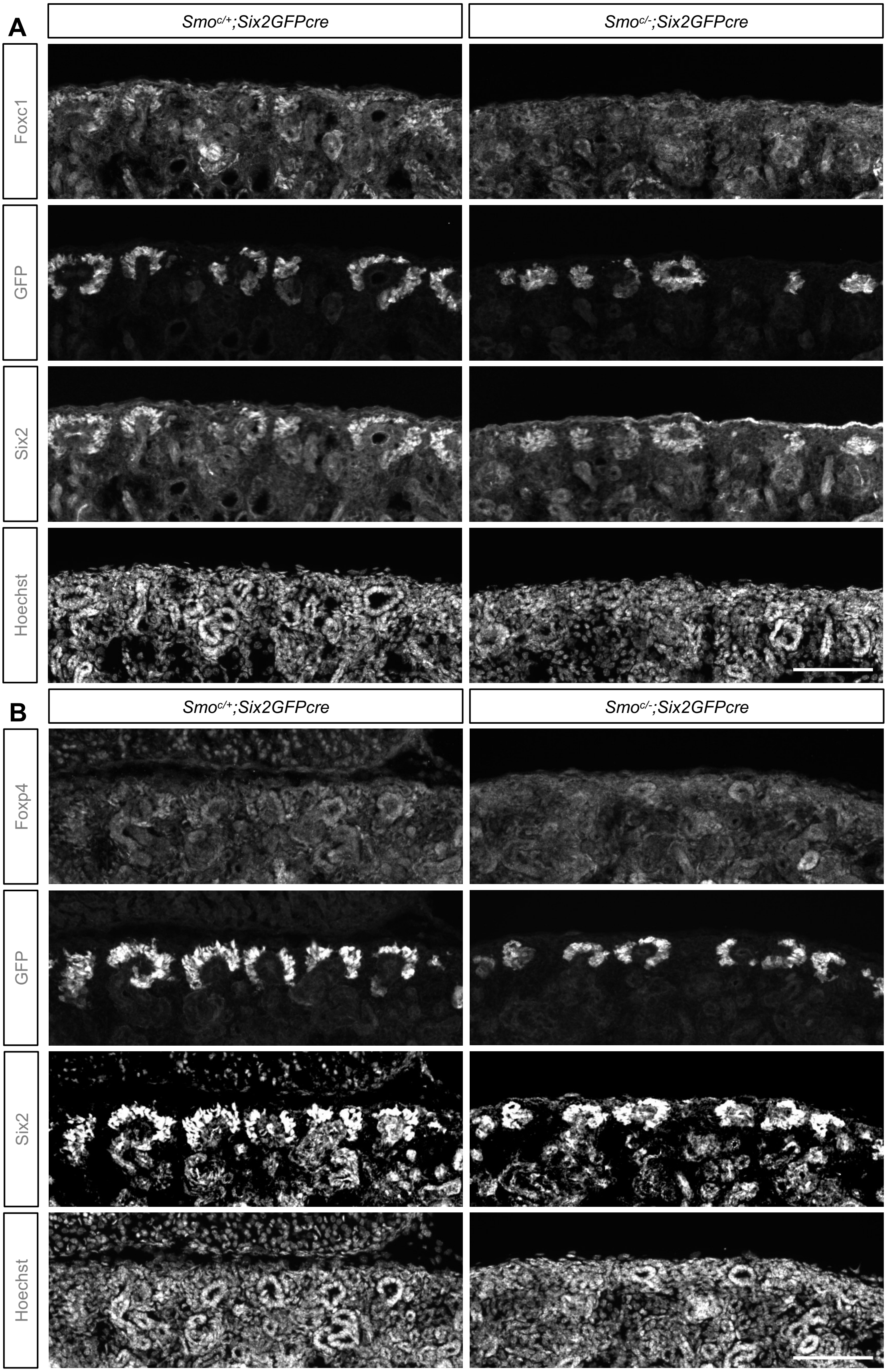


Supplemental Figure 6
Immunofluorescence analysis shows that mesenchymal nephron progenitors lacking Smo show weaker Foxc1 (A) and Foxp4 (B) signals. Foxc1 and Foxp4 are also detectable in podocytes and collecting duct, respectively. GFP represents Six2GFPcre, showing good overlap with endogenous Six2. Stage, E18.5. Scale bar, 100μm.

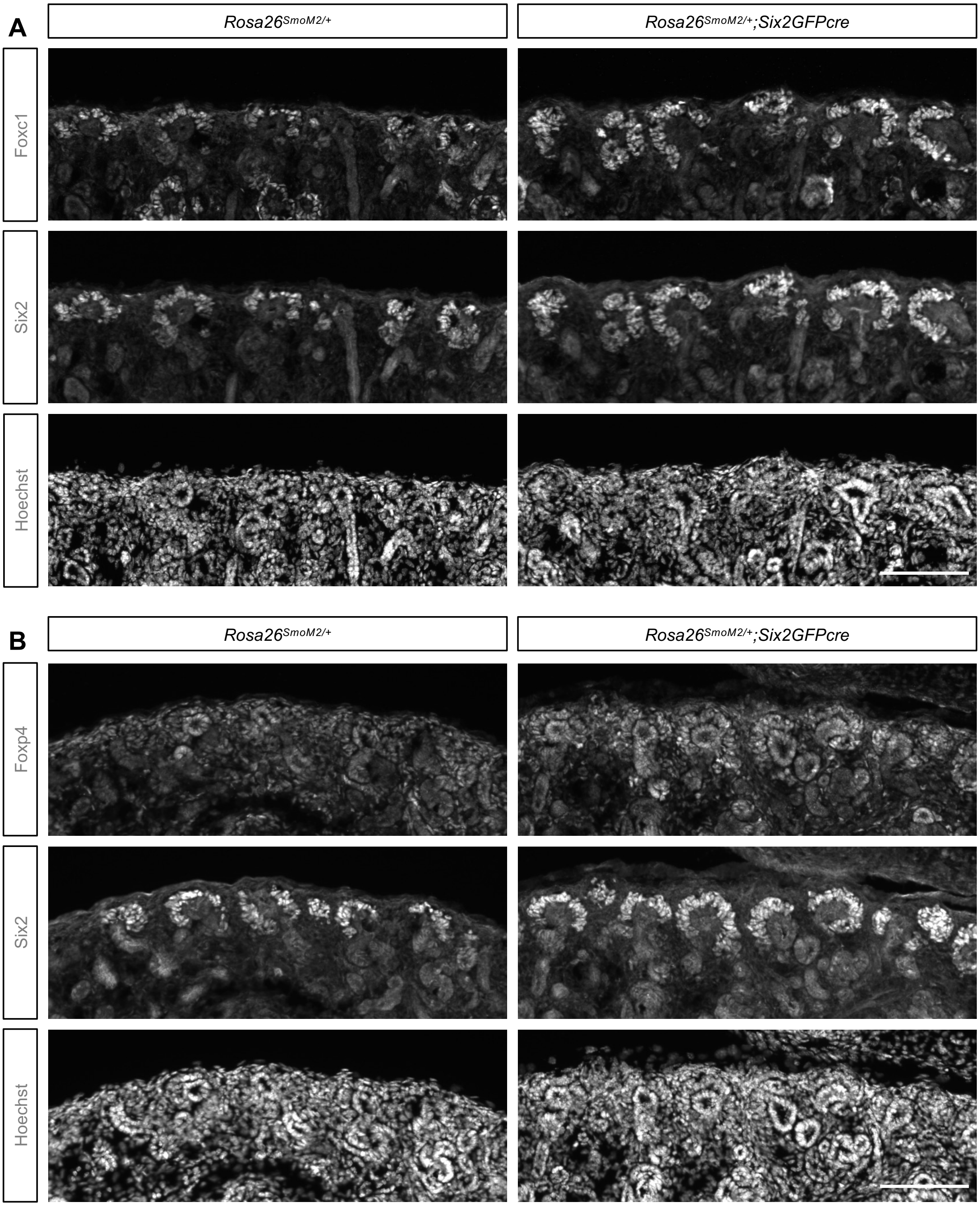


Supplemental Figure 7
Immunofluorescence analysis shows that mesenchymal nephron progenitors expressing SmoM2 show stronger signals of Foxc1 (A) and Foxp4 (B). Foxc1 and Foxp4 are also detectable in podocytes and collecting duct, respectively. Stage, E18.5. Scale bar, 100μm.

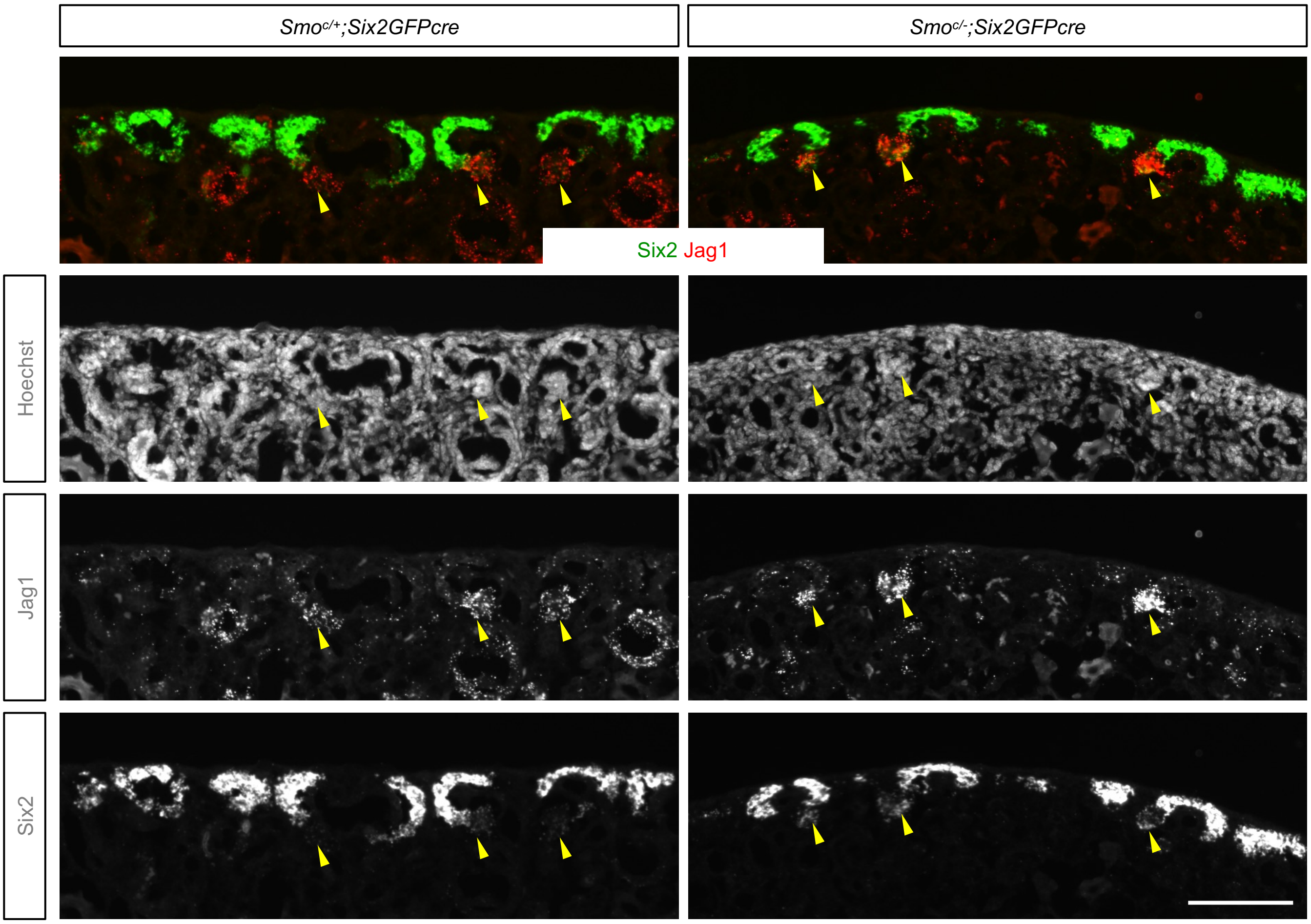


Supplemental Figure 8
RNAscope in situ hybridization for *Six2* and *Jag1* in the Smo loss-of-function mutant and control kidneys. Yellow arrowheads mark pretubular aggregates or renal vesicles where *Six2* is downregulated. In the Smo mutant kidney, activation of Jag1 is more robust. Given Six2 protein is detectable in pretubular aggregates and renal vesicles, it is likely that those cells expressing Jag1 were included in our FACS-isolated cells, contributing to apparent increase in *Jag1* expression detected in mNPs lacking Smo. Stage, E18.5. Scale bar, 100μm.

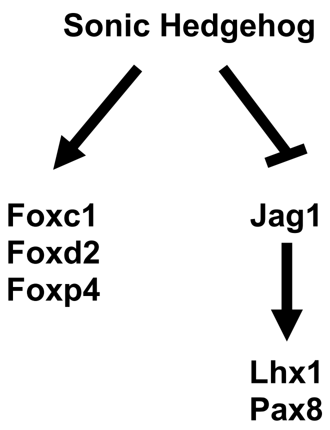


Supplemental Figure 9
Model of Shh action in mesenchymal nephron progenitors.
